# The pharmacodynamic inoculum effect from the perspective of bacterial population modeling

**DOI:** 10.1101/550368

**Authors:** Desiree Y. Baeder, Roland R. Regoes

## Abstract

The quantitative determination of the effects of antimicrobials is essential for our understanding of pharmacodynamics and for their rational clinical application. Common pharmacodynamic measures of antimicrobial efficacy, such as the MIC and the pharmacodynamic function, fail to capture the observed dependence of efficacy on the bacterial population size — a phenomenon called inoculum effect. We assessed the relationship between bacterial inoculum size and pharmacodynamic relationship and determined the consequences of the inoculum effect on bacterial population dynamics with a mathematical multi-hit model that explicitly describes the interaction between antimicrobial molecules with their targets on the bacterial cells. Our model showed that the inoculum effect can arise from the binding dynamics of antimicrobial molecules to bacterial targets alone. A pharmacodynamic function extended by the inoculum effect on its parameters was able to predict the long-term population dynamics of simple scenrios well. More complex scenarios, however, were only captured with by the mechanistically more explicit multi-hit model. In simulations with competing antimicrobial-susceptible and -resistant bacteria, neglecting the inoculum effect led to an overestimation of the competitive ability of the resistant strain. Our work underpins the importance of including the inoculum effect into quantitative pharmacodynamic frameworks, and provides approaches to accomplish that.

## Introduction

Efficacy of antimicrobials against bacteria is commonly quantified using time-kill curves. The slopes of a log-linear function fitted to time-kill curves (Figure 1a) describe the change of the bacterial population over time and are a direct measure of the bacterial net growth rate. The net growth rate and its dependence on the antimicrobial concentration can be captured by the so-called pharmacodynamic (PD) function.^1,2,3^ The parameters of this PD function, however, depend on the initial bacterial population size in the time-kill curve experiments.^4,5^

**Figure 1:**
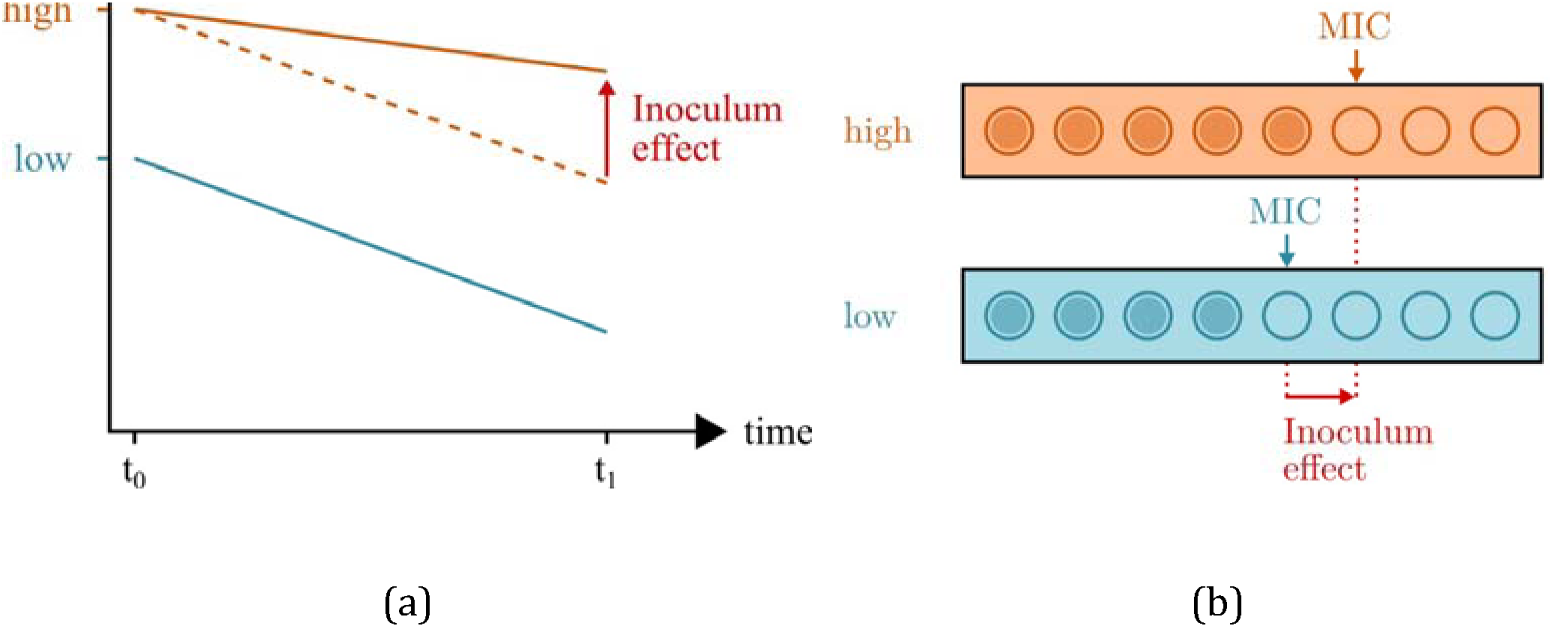
The inoculum effect changes efficacy of antimicrobials. Two common methods to quantify the efficacy of antimicrobials are (a) time-kill curves and (b) MIC measurements. (a) Time-kill curves illustrate the change in population size over time in presence of antimicrobials. The bacterial inoculum is the bacterial population size *N* at *t0 = 0h* (*N_inoc_ = N(t = 0)*). The slope of the curve results from fitting a log-linear function to the population dynamics. Here, only the log-linear curves and not individual measurements are shown. The slope of the time-kill curve representing dynamics of bacteria starting at a high inoculum size (orange) is shallower than dynamics of bacteria starting at a low inoculum (blue). In the absence of an inoculum effect, the slope is independent of the inoculum (dashed orange line). (b) Alternative to measuring time-kill curves, efficacy is typically determined with the MIC. An inoculum effect is observed if increasing the inoculum size from low (blue) to high (orange) increases the MIC.

A more widespread, but less informative, way to express efficacy of antimicrobials is the MIC (Figure 1b). Although it is recognized that the MIC might change with bacterial inoculum size (see for example the MIC protocol by Wiegand *et al*.^6^), both the CLSI and the EUCAST recommendations to measure the MIC are based on a single standard bacterial inoculum of 5 × 10^5^ cfu/mL.^7,8^ This standardization of the inoculum yields an estimate of the MIC that cannot be extrapolated to other bacterial population sizes.

Changes in efficacy of antibacterial drugs caused by changing the initial bacterial inoculum size are summarized under the term *inoculum effect* (Figure 1). The inoculum effect was first described in the early 1940s,^9^ and was observed for a variety of combinations of bacterial species and antimicrobials more recently.^4,10,11,12,13,14,15,16,17,18,19^ In addition to its experimental establishment, the inoculum effect has been explored with mathematical models.^4,13,19,20,21,22,23,24^ These experimental and theoretical studies have put forward hypotheses explaining the inoculum effect that fall into two broad categories. The first category attributes the inoculum effect to the binding kinetics of antimicrobials: the higher the bacterial inoculum the more antimicrobial molecules are bound, which leads to a larger reduction of free antimicrobial molecules.^4,21,22^ The second category invokes the secretion enzymes at high bacterial inocula, e.g. *β*-lactamases^9,10,14,15,16,25,26,27^ and proteases.^28,29,30^

In this study, we used mathematical models as a tool to investigate these competing hypotheses about the inoculum effect. Specifically, we use the so-called multi-hit model^31^ to address if the binding kinetics by itself already leads to an inoculum effect, and to what extent it influences the PD parameters. We have already used this model previously^32^ and a similar, albeit more antimicrobial-specific approach has been used to model classic reaction kinetics of antibiotics.^21^ Here we present an additional implementation of this model by exploring the effects of the bacterial inoculum conceptually. Additionally, we investigate how the inoculum effect biases the competition between antimicrobial-susceptible and -resistant strains.

## Methods

### Multi-hit model

We used the multi-hit model^21,31,32^ that describes a bacterial population consisting of *N_tot_* cells. The key idea of this model is that targets on bacterial cells are “hit”, i.e. bound, by antimicrobial molecules, and a bacterial cell dies if a threshold number of targets, which we call *z*, has been hit. The total number of hits possible per bacterial cell is called *n* Antimicrobial molecules attach to and detach from a bacterial targets with the rates *α_i_* and *µ_i_*, respectively. We denote the number of bacteria that have i antimicrobial molecules adsorbed to cellular targets by N_i_. Bacteria die with Bacteria die with a death rate *d_i_*. Formally, the bacterial population in the multi-hit model is structured according to how many antimicrobials have hit the cell (Figure 2). For details of the multi-hit model, see supplement S1.

**Figure 2:**
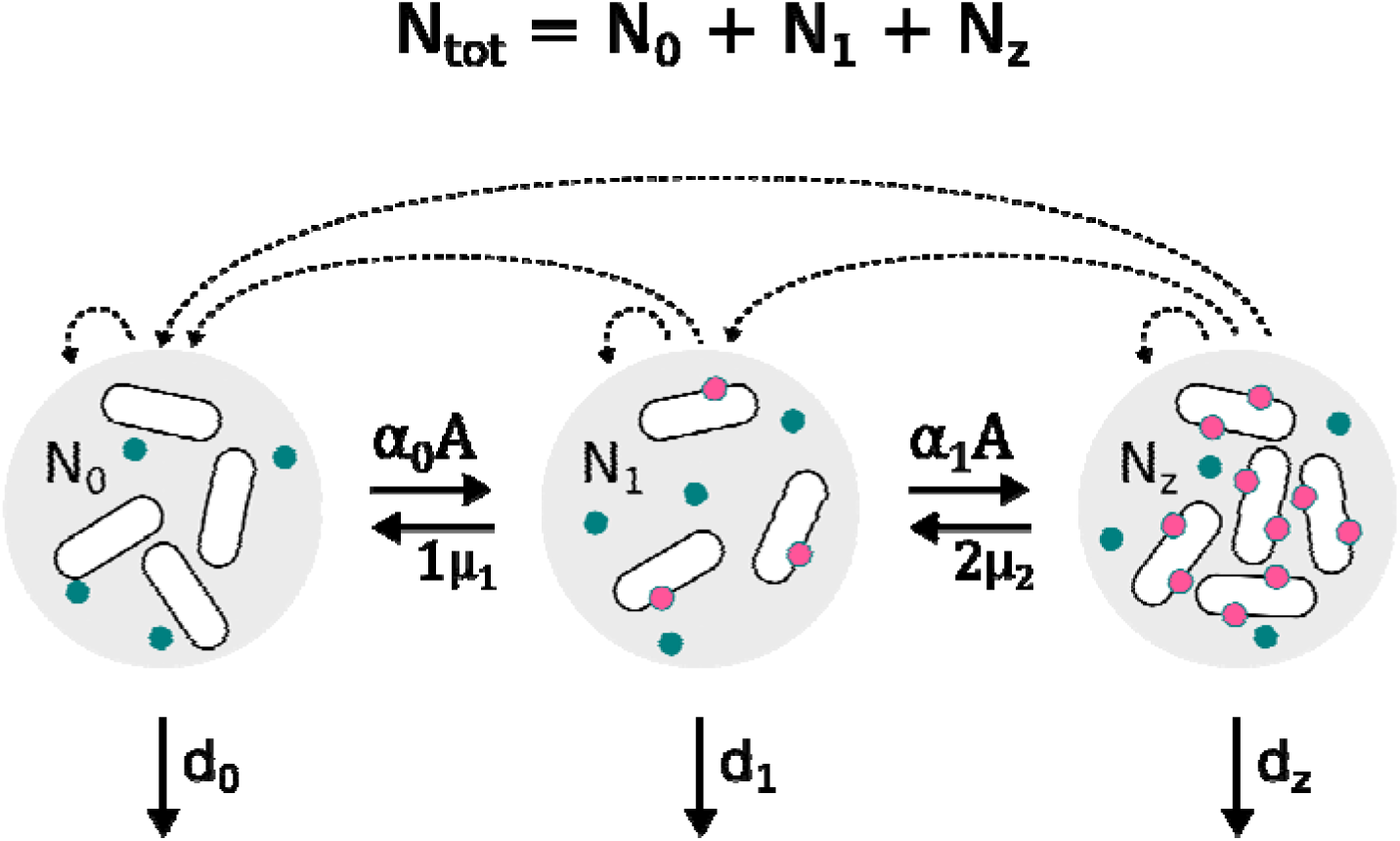
Schematic representation of the basic multi-hit model with three classes (*n* = 2). At every time point the bacterialgains bacterial cells from the population *N_tot_* can be divided in *n* subpopulations based on the number of antimicrobials that have hit the bacterial cells. A hit is the mathematical simplification of adsorption and subsequent downstream processes connected to binding of one antimicrobial molecule to a bacterial target site and is described with the rate *α*. The parameter *µ* is the detachment of an antimicrobial from the bacterial target. We assumed that finding the bacterial cell is the rate limiting step for an antimicrobial molecule.^59^ This is appropriate if the number of targets are in abundance, e.g. when antimicrobial peptides adsorb to the cell membrane of bacterial cells. For antimicrobials that act within the cell, a hit rate dependent on the number of free targets is more appropriate (see^21^). Each class *N_i_* gains bacterial cells from the class *N_i_*_−1_ with the term *α_i_*_−1_*AN_i_*_−1_ and a bacterium dies with an increased death rate *d_z_* if two antimicrobial molecules have hit the cell, i.e. *z* = 2. Therefore, *N_tot_* = *N*_0_ + *N*_1_ + *N*_2_, with *N*_2_ = *N_z_*. *A* is the number of free antimicrobial molecules, here indicated in blue, antimicrobials that have hit a cell are indicated in pink. The rates of the ODE (see supplement S1 for equation) are indicated with arrows and bacterial replication with dashed arrows.

### Variations in the Multi-hit framework

In the basic model (equation 1–3 in supplement S1), we assumed the case of complete multihit,^31^ with *z* = *n*, which maybe best describes antimicrobials that need to hit a large fraction of the existing targets, for example AMPs.^33^ Moreover we assumed that the total amount of antimicrobials is stable and that antimicrobial molecules are recycled back into the system when a cell dies.

We considered variants of the multi-hit model by relaxing some of the assumptions above: (i) not all targets need to be hit *z* < *n*, which leads to non-effective binding, (ii) non-reversible binding, (iii) unspecific binding (i.e. binding to non-targets) in combination with non-reversible binding, leading to molecules attaching non-reversibly to targets and non-targets alike, and (iv) enzymatic degradation. For details and model equations, see Supplement S2.

### Pharmacodynamic and Pharmacokinetic function

To investigate how the efficacy of antimicrobials changes with the inoculum in our multi-hit model simulations, we used the PD function^1,2,3^ as a measure of antimicrobial efficacy. Specifically, we calculated the bacterial net growth rate *ψ* on basis of the antimicrobial concentration *A* at the start of the simulation (*A*(*t* = 0) = *A*_0_), with

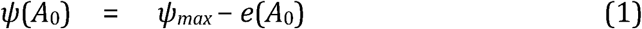

In this common conceptualization of PD effects, the net growth rate *ψ* is divided into the growth rate of the bacteria in absence of the antimicrobials *ψ_max_* and an effect *e*(*A*_0_) that depends on the concentration of the antimicrobials. Note that we use the concentration *A*_0_, because throughout the simulation, A varies due to binding and degradation.

The effect *e*(*A*_0_) is further parameterized by the concentration, *MIC_PD_*, that leads to a zero net growth rate, *ψ*(*A*_0_ = *MIC_PD_*) = 0, the minimum net growth rate that can be obtained with antimicrobials *ψ_min_* = *ψ*(*A*_0_ ⍰∞), and a parameter that characterizes the steepness of the pharmacodynamic relationship, *κ*:

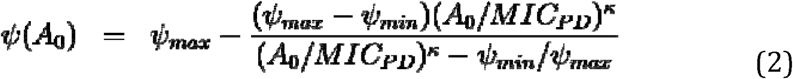

Here, the *MIC_PD_* approximates the *MIC* if the antimicrobial is concentration independent.^34^

To estimate the PD parameters, we simulated time-kill curves for which we varied the antimicrobial concentration (Figure S1). In a first step, the net growth rate was calculated based on the simulated time-kill curves using the relationship:

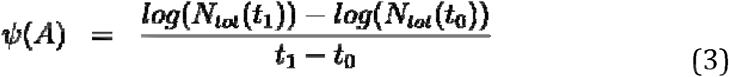

The time window, t_1_-t_0_, is typically chosen to be on the order of hours^1,32,35^ **Error! Reference source not found**.,^36^ to allow the measurement of potentially steep declines of the bacterial population size at high antimicrobial concentrations. For example, Tobramycin decreases the bacterial population faster at high concentrations than Ticarcillin^35^. Thus, to estimate the PD parameters, a shorter time window needs to be chosen for Tobramycin than for Ticarcillin. Additionally, for wider time windows, more complexities arise, such as persister populations that are detectible within a few hours.^37,38,39,40^ For the sake of simplicity, we set the time window that we use to calculate the growth or decline of the bacterial population in our simulations to 1 hour. The time window can be set to be narrower or wider when, instead of aiming for a generic understanding of the inoculum effect, a specific bug-drug combinations is being investigated.

The determined net growth rates were then used to estimate the PD parameters. The steepness parameter *κ* was calculated (see supplement S3). The estimates of the PD function parameters differed by inoculum. This dependence of the PD parameters were captured by an extended PD function, *ψ*(*A*_0_,*N_inoc_*). Specifically, the inoculum effect was quantified as slope *m* of the function

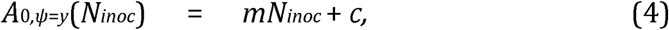

with *y* = 0 and *y* = *E_max_*/2 in the case of *ψ*(*A*_0_ = *MIC_PD_*) and *ψ*(*A*_0_ = *A*_50_), respectively.

Note that the pharmacodynamic function does not capture evolutionary processes. However, evolved bacterial strains can be incorporated into this pharmacodynamic modeling framework as subpopulations with their own set of PD parameters.^36,41^

The pharmacokinetic function we used is described in Regoes *et al*.^1^ In short, the initial antimicrobial concentration *A*_0_ is assumed to decay with a rate *k_A_*, and reset to *A*_0_ every 8 hours.

### Population model

To investigate the agreement between the bacterial population dynamics predicted by the common parameterization of antimicrobial efficacy with PD functions and those predicted by the more detailed multi-hit model, we incorporated the extended PD function and the pharmacokinetic function into a population model:

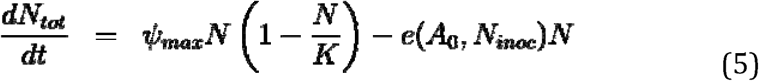

Here, *K* is the carrying capacity. We used this framework before in.^42^

### Competition

To describe competition between an antimicrobial-susceptible and an antimicrobial resistant strain, we extended the basic multi-hit model with a second strain. The number of susceptible and resistant cells are denoted by *S* and *R*, respectively. In our simulations, strain *R* is assumed to have a 10 times lower attachment rate, which could correspond to surface modification on resistant bacterial cells — a common resistance mechanism against AMPs.^28^ For more details and model equations, see Supplement S4.

### Implementation

The multi-hit model was implemented in R^43^ (www.r-project.org^;^ version 3.4.0). We solved the deterministic multi-hit model numerically with the package deSolve.^44^ For plotting, we used the following packages: sfsmisc^45^ and RColorBrewer.^46^ The code is available upon request.

## Results

We developed a mathematical model to describe the interaction between a bacterial population and antimicrobials. The model is based on the multi-hit concept of antimicrobial action,^31^ according to which bacteria are killed as soon as a certain number of antimicrobial molecules hit the cell. This modeling framework describes the interaction between antimicrobial molecules and bacterial cells in more detail than common population biological models, and thus allowed us to investigate the determinants of the inoculum effects and predict quantitatively how the inoculum affects the pharmacodynamics.

### Binding kinetics alone generates an inoculum effect

First, we investigated if an inoculum effect emerges simply as a result of concentration-dependent interactions between bacteria and antimicrobials and their binding kinetics. To that end, we used the basic multi-hit model that assumes reversible binding of antimicrobial molecules to target sites on the bacterial cells.

We found that an inoculum effect emerged. In our simulations, a bacterial population starting at a high inoculum grew over time, while a smaller initial population exposed to the same amount of antimicrobials declined (Figure 3a). The explanation of the effect can be found in the concentration of free antimicrobial molecules that is sensitively affected by the bacterial inoculum (Figure 3b), resulting in a lower efficacy if the inoculum is high. Consistent with this explanation, the inoculum effect was abrogated if the free antimicrobial concentration was assumed to be constant independent of any binding processes (Figure S2). Thus, only the binding of antimicrobial molecules — even though it was assumed reversible — caused an inoculum effect. Allowing for degradation of antimicrobials by extracellular enzymes in our model exacerbated the inoculum effect (Figure 3) but was not necessary.

**Figure 3:**
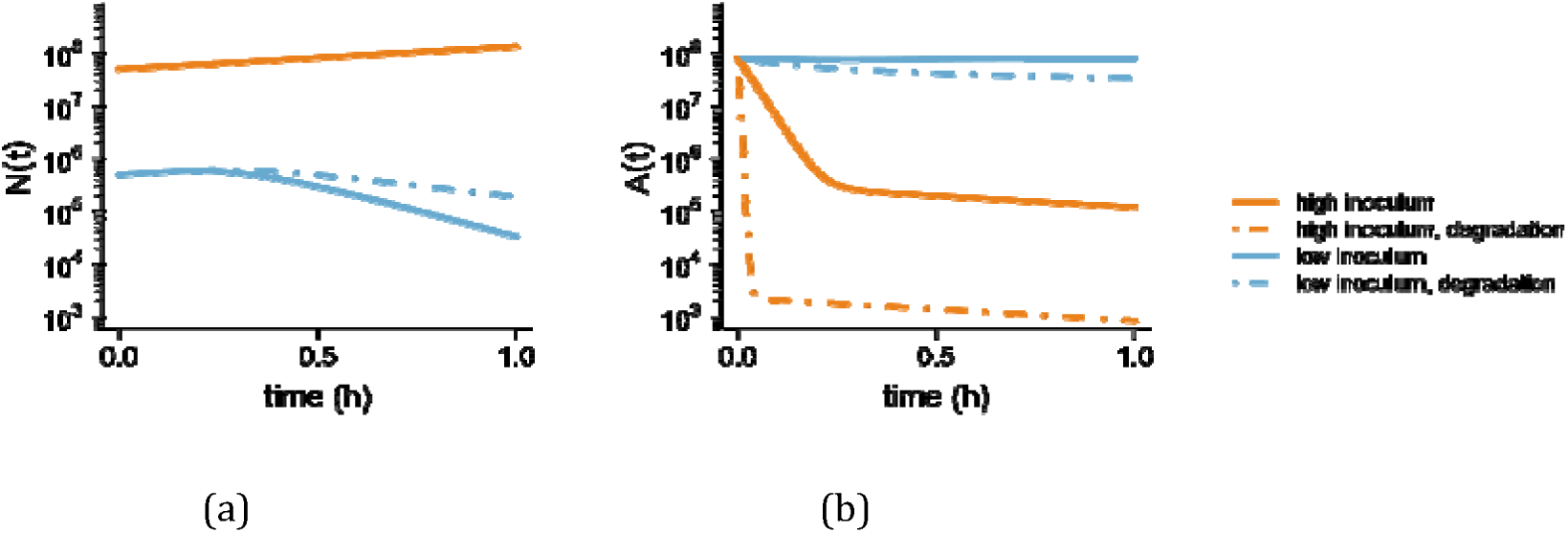
Increasing the bacterial inoculum size from 5 ⍰ 10^5^ (blue) to 5 ⍰ 10^7^ (orange) changes the efficacy of the antimicrobial due to reduction of the number of free antimicrobials. Simulations with reversible binding is shown with solid lines, dashed lines indicate simulations with degradation. (a) The bacterial inoculum *N(t)* is affected differently by antimicrobials depending on bacterial inoculum size and binding (b). Time course of free antimicrobial molecules. The range of the decline of free antimicrobials is consistent with other modelling studies^19^ , and *in vitro*^33^ and *in vivo* observations^35^. Parameters not mentioned here are listed in Table S1.

### Inhibitory concentrations increase linearly with inoculum, but with a proportionality constant < 1

This inoculum effect can be quantitatively captured. Specifically, we studied how the inoculum affects the parameters of the *pharmacodynamic function* We considered the effect of the inoculum on the pharmacodynamic function that describes relationship between the net growth rate of bacterial population and the antimicrobial concentration (see Methods). Performing *in silico* time kill experiments, we estimated the pharmacodynamic functions for different inocula (Figure 4a) and found that the efficacy across all antimicrobial concentrations is lower for larger inocula.

**Figure 4:**
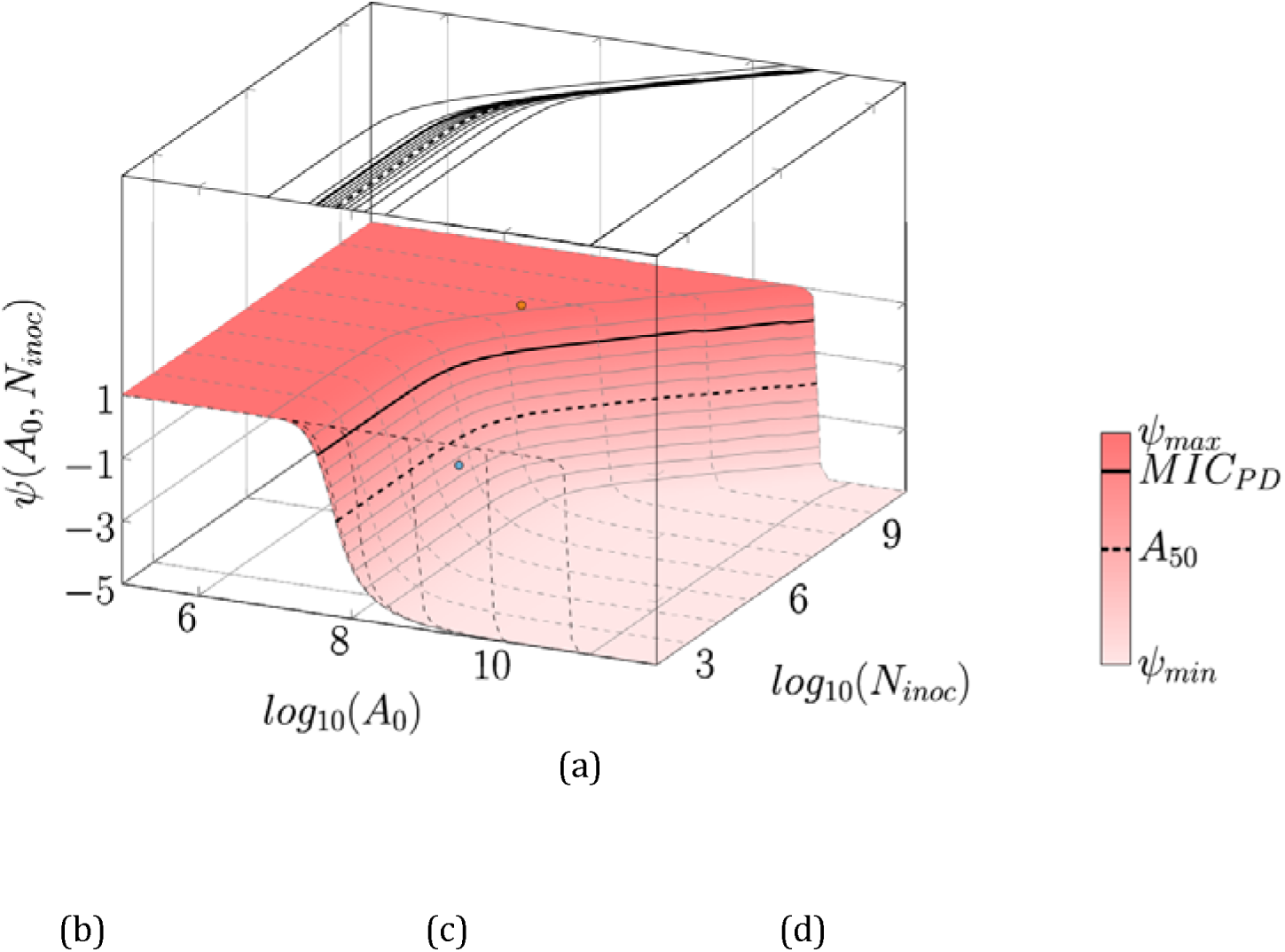
Inoculum size affects pharmacodynamics. (a) We determined the efficacy for varying inoculum sizes (N_inoc_) with z = 10, resulting in a PD plane (note that in contrast to (b)-(d), the inoculum effect here is illustrated on a log_10_-log_10_ scale). In the PD plane, solid gray contour lines show combinations of antimicrobials and inoculum sizes with the same net growth rate, both in the PD plane and projected above the PD plot. *MIC_PD_* and *A*_50_ are indicated as bold solid and dashed line, respectively. For several inoculum sizes, we plotted the PD curve as dashed contour lines along the antimicrobial molecule number axis and again projected the contour lines, this time in front of the plot. The slopes of Fig. 3a (solid lines). are marked as dots in the corresponding colors. In (b)-(d) we plotted the PD parameters *MIC_PD_*, *A*_50_, and κ against the bacterial inoculum for varying number of hits needed to kill a bacterial cell (z). The bold black line in (b) corresponds to the bold solid line in (a) and the bold black line in (c) correspond the bold dashed lines in (a). Parameters not mentioned here are listed in Table S1.

The most important efficacy parameter, the pharmacodynamic parameter *MIC_PD_* increases linearly with the inoculum (Figure 4b). However, the proportionality constant is less than one, meaning that a doubling of the inoculum does not lead to a doubling of number of antimicrobial molecules required to halt bacterial population from growing, and thus does not lead to a doubling of the *MIC_PD_*. We found a similar linear increase of another common efficacy measure, the *A*_50_ (see Methods for definition and Figure 4c), which is interestingly also linearly affected by *z*.^32^ We also found that these two efficacy parameters increased linearly with the number of hits required to kill a bacterial cell, *z* (Figure 4b,c) and that the increase was steeper when we allowed for degradation (Figure S3). Furthermore, the steepness parameter κ did not increase linearly, but saturated (Figure 4d). These results hold for different modes of binding (Figure S4).

Formally, we found the following relationship between the efficacy parameters and the inoculum *N_inoc_*:

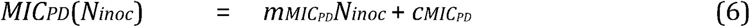

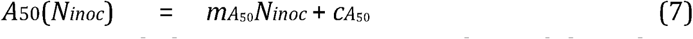

Estimates for the intercept and slope parameters, c and m, of these linear relationships are shown in Table 1. With these two parameters, we can also describe how many more antimicrobial molecules we need to achieve the same pharmacodynamic efficacy when we increase the bacterial inoculum *x*-fold:

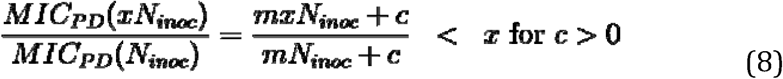

**Table 1:**
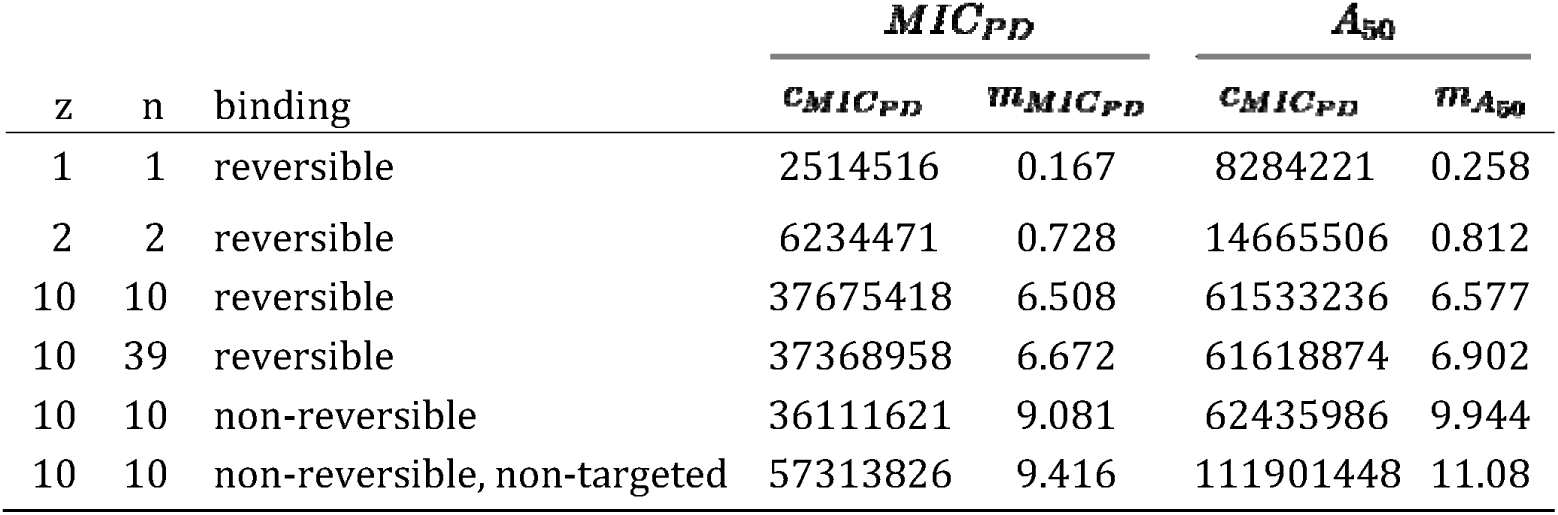
Results of fitting linear function *x* = *m_x_N_inoc_* +*c_x_* with *x* = **MIC_PD_** and *x* = *A*_50_ to the simulated data. Reversible binding with *z* = 1,2,10 is also plotted in Figure 4b-d.

|  |  |  | *MIC<sub>PD</sub>* |  | *A<sub>50</sub>* |  |
| --- | --- | --- | --- | --- | --- | --- |
| z | n | binding | *c<sub>MIC<sub>PD</sub></sub>* | *m<sub>MIC<sub>PD</sub></sub>* | *c<sub>A<sub>50</sub></sub>* | *m<sub>A<sub>50</sub></sub>* |
| 1 | 1 | reversible | 2514516 | 0.167 | 8284221 | 0.258 |
| 2 | 2 | reversible | 6234471 | 0.728 | 14665506 | 0.812 |
| 10 | 10 | reversible | 37675418 | 6.508 | 61533236 | 6.577 |
| 10 | 39 | reversible | 37368958 | 6.672 | 61618874 | 6.902 |
| 10 | 10 | non-reversible | 36111621 | 9.081 | 62435986 | 9.944 |
| 10 | 10 | non-reversible, non-targeted | 57313826 | 9.416 | 111901448 | 11.08 |

The same relationship holds for the *A*_50_, but, of course, with *A*_50_-specific estimates for *c* and *m*. Thus, the increase in antimicrobials required to achieve the same effect on an x-fold increased inoculum is, according to our model, less than x-fold and thus the proportionality constant is less than one.

The parameters characterizing the net bacterial growth rate in absence of antimicrobials is independent of the inoculum (Figure S5a). Also, the maximum effect that can be achieved with an antimicrobial, *E_max_*, and the related minimum net growth rate, *ψ_min_*, are independent of the inoculum (Figure S5b-c). This is a consequence of the fact that these parameters are not affected by the binding of antimicrobial molecules to targets on the bacterial cells.^32^

The PD function *ψ*(A0) has been used to predict the long-term bacterial population dynamics from antimicrobial effect on the initial net growth rate.^1,4,5,35,42,47,48,49^ However, if we include the inoculum effects on *MIC_PD_*, A50, and κ into the PD function, we found that this extended PD function therefore captured only the short-term bacterial population dynamics while only the mechanism-based multi-hit model could describe more complex population dynamics (see supplement S5, Figure S6, Figure S7).

### Inoculum effect alters the outcome of bacterial competition

To shed light on the effects of the inoculum on the competition between bacterial strains, we extended the multi-hit model framework by an additional resistant strain *R*. We assume that strain *R* carries a fitness cost and is hence not able to outcompete strain *S* at low antimicrobial concentrations (Figure 5a). At intermediate concentrations, *R* outcompetes strain *S* (Figure 5b). Lastly, at high concentrations, both strains are killed (Figure 5c). The antimicrobial concentrations that mark the transitions between these three dynamical outcomes define the so-called *mutant selection window* (MSW).^50,51^

**Figure 5:**
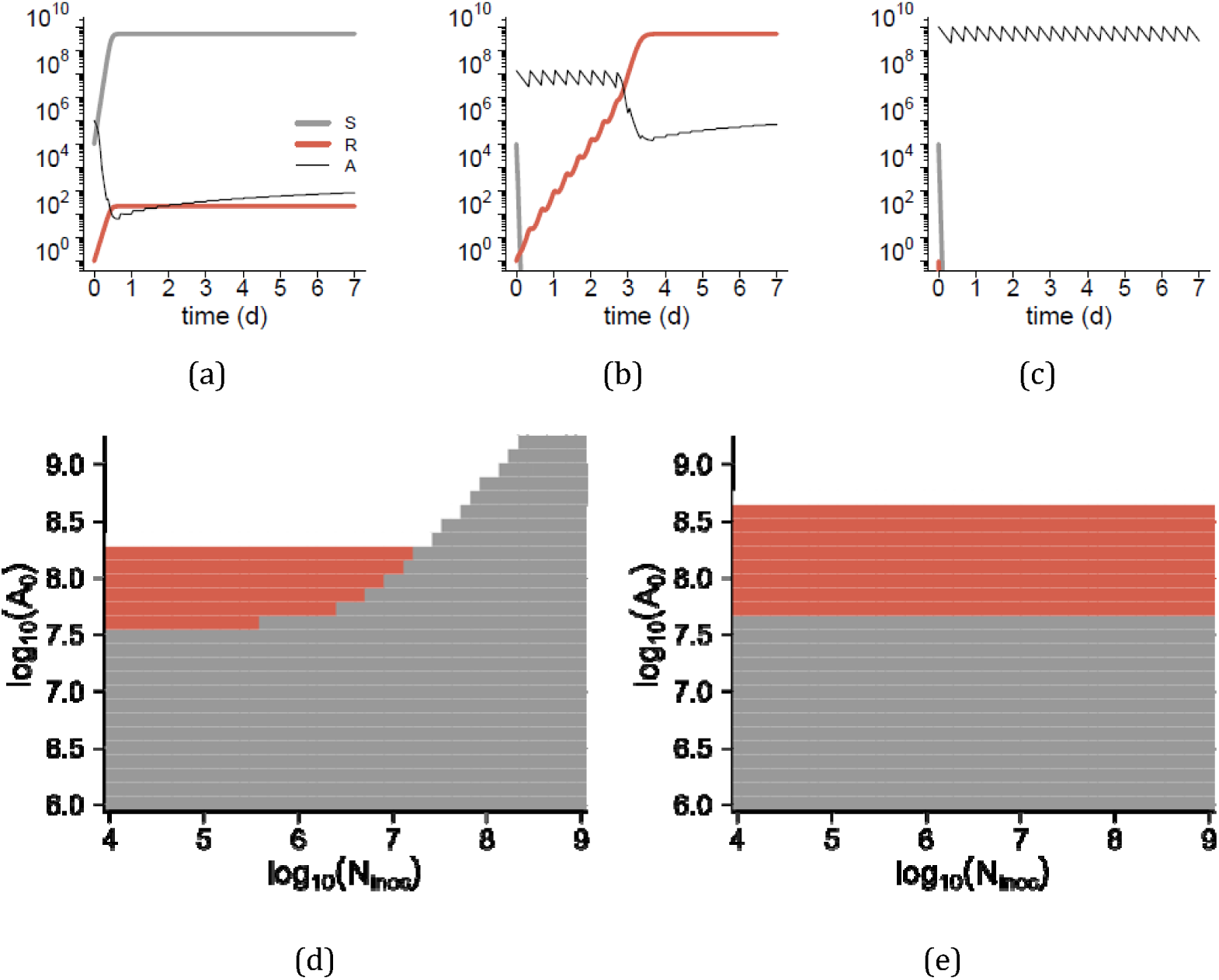
The inoculum effect influenced competition outcomes. Antimicrobial doses *A*_0_ changed the outcome of competition: (a) *S* won when exposed to low antimicrobial doses, (b) *R* won at intermediate antimicrobial doses and (c) bacteria went extinct at high antimicrobial doses. (d) In an additional dimension, the inoculum effect changed the outcome of competition (gray: *S* is more abundant, red: *R* is more abundant, white: extinction (*S,R <* 1) at the end of the simulation). (e) For comparison, we used the PD function to simulate competition without inoculum effect.

To investigate how the inoculum affects the size of the MSW, we conducted simulations across a range of inocula. In these simulations we also included a time frame of 7 days and repeated doses of the antimicrobial, to mimic antimicrobial usage in a treatment scenario. We therefore use the term” treatment MSW”. The difference between the MSW and the treatment MSW is that the MSW is based on time-kill curves with a single input of antimicrobials, while the treatment MSW is based on time-kill curves in treatment regimes.

We find that the treatment MSW shrinks and even vanishes for increasing inocula (Figure 5d). Qualitatively similar results hold for simulations based on the extended PD function (Figure S8). Simulating bacterial competition with a model that does not capture the inoculum effect, in contrast, left the MSW unaffected (Figure 5e).

## Discussion

In this study, we used a mathematical model to understand how an inoculum effect can arise and to investigate how the bacterial inoculum affects the pharmacodynamics of antimicrobials. With our model simulations, we were able to derive predictions for the inoculum effect not only on the MIC, but on multiple pharmacodynamic parameters. While the insights from our study are generic and conceptual, they can be adapted to any specific antimicrobial-bacterial pathogen combination. Our predictions give rise to many experimentally testable hypotheses that we detail below.

Udekwu *et al*.^4^ previously showed how the inoculum effect on the MIC can be integrated into the PD function. We extended these efforts to the additional pharmacodynamic parameters: the antibiotic concentration at which 50% of the maximum effect is obtained, *A*_50_, and the parameter that describes the steepness of the pharmacodynamic relationship, *κ*. We found that the inoculum affected the PD parameters *MIC_PD_* and *A*_50_ linearly, and the parameter *κ* non-linearly, resulting in an inoculum effect that does not simply scale with the number of bacterial cells and antimicrobial molecules in the system. For example, to obtain the same effect on a, say, 100-fold increased bacterial population, we require less than 100-fold increase in the number antimicrobial molecules. This finding directly contradicts the argument that the inoculum effect is an artifact^27^ arising from simple scaling. We could show that to achieve the same antimicrobial effect for a higher inoculum always requires a smaller increase in the number of antimicrobial molecules than a simple scaling would suggest. How much smaller depends on the specific parameters of the bacteria, antimicrobials, and assay system. Interestingly, this means that, on a per-molecule level, antimicrobials become more effective with increasing inoculum.

The mechanistic underpinning of the inoculum effect in our simulations is that an increase in the bacterial inoculum results in higher depletion of free antimicrobials. We call this the *numbers game*: Binding, even if it is reversible, takes molecules out of the pool of free antimicrobial molecules, which in turn means that there are fewer molecules available for binding in the next time step. For larger bacterial inocula, the antimicrobial depletion by binding is increased, resulting in the inoculum effect. Thus, for a given antimicrobial-bacterial pathogen combination, we expect an inoculum effect if the numbers game is at play. This numbers game results in an inoculum effect has already been observed in mathematical modeling studies.^21,31^ Experimental studies that investigate the possibility and extent of an inoculum effect arising from the numbers game alone^4,18,21^, however, are rare. Often, the inoculum effect is measured in systems in which additional effects, e.g. non-targeted binding^52^ or enzymatic degradation^10,11,12,14,15,16,25,53,54,55^, occur. These effects strongly increased the complexity of the population dynamics in our simulations. We therefore suggest conducting experimental studies that address the inoculum effect first in systems that are governed only by a simple binding kinetics of antimicrobials to their targets. These experiments should be conducted in a regime of declining free antimicrobial molecule concentrations with increasing bacterial inoculum sizes, because, in the numbers game scenario, this decline causes the inoculum effect. Such studies will form the basis of a quantitative understanding of the inoculum effect and will provide valuable baselines for studies of systems with more complexity.

In addition to determining if the binding kinetics of antimicrobials alone results in an inoculum effect, we suggest experiments that assess the more qualitative prediction of the multi-hit modeling framework we presented here. For example, we showed that the more targets are needed to be hit to kill a cell (*z*), the more pronounced the inoculum affect was. In the case of antimicrobial peptides, z varies from 80-800 in the case of protegrin^56^ to 10^6^ −10^7^ in the case of PMAP-23.^33^ With everything else being equal, we therefore expect a more pronounced inoculum effect for PMAP-23 than for protegrin. Similar predictions can be easily derived for other antimicrobials if quantitative parameters characterizing their mechanisms of action are known. Generally, the quantitative relationships between the mechanisms of action of an antimicrobial, its pharmacodynamics, and the size of the inoculum effect represent a promising area of future theoretical and experimental studies.

The inoculum effect has been demonstrated experimentally for a number of bacterial strains and antimicrobials. ^4,10,11,12,14,15,16,17,18,19,21,53,54,57^ In most cases, the MIC was used to determine and quantify the inoculum effect.^4,10,11,12,14,17^ Our comprehensive analysis of the inoculum’s effect on all pharmacodynamic parameters showed that the steepness parameter κ increases. Therefore, the inoculum effect is stronger for *A*_50_ than for *MIC_PD_* and determining the effect on *A*_50_ represent the most sensitive measure of the inoculum effect. Based on this observation, we recommend to not only quantify the *MIC*, but the whole pharmacodynamic relationship, and to use OD measurements with caution because they do not allow the estimation of the pharmacodynamic relationship for antibiotic concentrations above the *MIC_PD_*.

The absence of the inoculum effect has been reported repeatedly.^27,57^ Our results shed light on the conditions under which an inoculum effect is expected to arise. First, in our simulations the inoculum effect did not manifest itself in certain regions of the PD plane: for very small and very high antimicrobial molecule numbers, the inoculum effect was absent. Second, in our simulations, the inoculum effect essentially depends on a decline in free antimicrobial concentrations. Therefore, we expect the inoculum effect not to arise in any system, in which the antimicrobial concentration does not decline with increasing bacterial inoculum. Third, the establishment of the inoculum effect on the MIC is impaired because of the uncertainties arising from the doubling of the antimicrobial concentration that is required according to the standard protocol for MIC determination. To give a specific example: according to the standard protocol, “real” values of the MIC of 9 and 15 *mg/L* would be both estimated as 16 *mg/L*; therefore, an inoculum effects increasing the MIC from 9 to 15 *mg/L* would not be detected. Lastly, in many studies an arbitrary threshold of an 8-fold increase of the MIC for a 100-fold higher bacterial inoculum is used to define the inoculum effect^6,10,14,15,16^ clouding inoculum effects of smaller extent. Rather than setting such a cutoff, we suggest determining the *MIC_PD_* or, even better, the *A*_50_ from adequately powered time-kill experiments.

Most surprisingly, we found that the inoculum effect can have unexpected and wide-ranging consequences for the evolutionary dynamics of bacterial populations under drug treatment. Our simulation of bacterial competition between drug-susceptible and -resistant strains revealed that the competitiveness of drug-resistant strain is overestimated when the inoculum effect is not considered. We found that, in our simulations, the “treatment MSW” — an estimation of the MSW^50,51^ under drug treatment — shrinks for higher bacterial inoculum sizes. Although the treatment MSW is an idealized concept, that does not capture all complexities of bacterial evolution and competition, a negative effect of higher inocula on the emergence of resistant strains should be observable in competition experiments. In competition experiments between drug-susceptible and - resistant bacterial strains, such as those presented in e.g.^51^ , our analysis predicts the selection coefficients to decrease with inoculum. Hence the inoculum effect is generally expected to impact bacterial survival and evolution *in vitro*, within infected hosts, and potentially even on the epidemiological level.

## Acknowledgments

We thank Andrew Read for sparking our interest in the topic of inoculum effects, Pia Abel zur Wiesch, Sebastian Bonhoeffer, and Jens Rolff for reading the manuscript, Bruce Levin for giving us valuable feedback on our discussion, and João Pires for interesting discussions about the project. This work was supported by an ETH Grant (ETH-41 15-2, R.R.R.) and by the Volkswagen-Stiftung (Az 96 695, R.R.R.).

